# Interactions between calmodulin and neurogranin govern the dynamics of CaMKII as a leaky integrator

**DOI:** 10.1101/809905

**Authors:** Mariam Ordyan, Tom Bartol, Mary Kennedy, Padmini Rangamani, Terrence Sejnowski

## Abstract

Calmodulin-dependent kinase II (CaMKII) has long been known to play an important role in learning and memory as well as long term potentiation (LTP). More recently it has been suggested that it might be involved in the time averaging of synaptic signals, which can then lead to the high precision of information stored at a single synapse. However, the role of the scaffolding molecule, neurogranin (Ng), in governing the dynamics of CaMKII is not yet fully understood. In this work, we adopt a rule-based modeling approach through the Monte Carlo method to study the effect of *Ca*^2+^ signals on the dynamics of CaMKII phosphorylation in the postsynaptic density (PSD). Calcium surges are observed in synaptic spines during an EPSP and back-propagating action potential due to the opening of NMDA receptors and voltage dependent calcium channels. We study the differences between the dynamics of phosphorylation of CaMKII monomers and dodecameric holoenzymes. The scaffolding molecule Ng, when present in significant concentration, limits the availability of free calmodulin (CaM), the protein which activates CaMKII in the presence of calcium. We show that it plays an important modulatory role in CaMKII phosphorylation following a surge of high calcium concentration. We find a non-intuitive dependence of this effect on CaM concentration that results from the different affinities of CaM for CaMKII depending on the number of calcium ions bound to the former. It has been shown previously that in the absence of phosphatase CaMKII monomers integrate over *Ca*^2+^ signals of certain frequencies through autophosphorylation (Pepke et al, Plos Comp. Bio., 2010). We also study the effect of multiple calcium spikes on CaMKII holoenzyme autophosphorylation, and show that in the presence of phosphatase CaMKII behaves as a leaky integrator of calcium signals, a result that has been recently observed *in vivo*. Our models predict that the parameters of this leaky integrator are finely tuned through the interactions of Ng, CaM, CaMKII, and PP1. This is a possible mechanism to precisely control the sensitivity of synapses to calcium signals.

## 1 Introduction

Information is stored in the brain through synaptic plasticity. It has been reported that synaptic strength is highly correlated with the size of the spine head, and the precision of information stored at a single synapse is quite high despite the stochastic variability of synaptic activation [6, 65, 61]. Structural changes to the postsynaptic spine that can lead to spine enlargement, and thus structural plasticity are triggered by *Ca*^2+^ signaling [39, 64]. Time averaging of these calcium signals has been suggested as a plausible mechanism for achieving the high precision of information processing observed in spines. Furthermore, phosphorylation of calcium/calmodulin-dependent protein kinase II (CaMKII) has been postulated as the most probable pathway satisfying the long time scales predicted for averaging [6].

CaMKII is an autophosphorylating kinase; in postsynaptic densities (PSD), CaMKII has been shown to play an important role in learning and memory [54]. Specifically, mice with a mutation in a subtype of CaMKII exhibit deficiencies in long-term potentiation (LTP) and spatial learning [86, 87]. Moreover, CaMKII expression regulates the rate of dendritic arborization and the branch dynamics [104, 97], highlighting its importance in structural plasticity. Additionally, CaMKII has been shown to bind actin, the major cytoskeletal protein in dendritic spines [14, 68], further emphasizing its role in structural plasticity.

Activation of CaMKII is exquisitely regulated at multiple levels as summarized below.

- **CaMKII activation by calmodulin (CaM)**: CaMKII is activated by calmodulin (CaM) [9], which is a protein with 4 *Ca*^2+^ binding sites: 2 on its C-terminal and 2 on N-terminal (Figure 1A) [98, 29]. CaM binds *Ca*^2+^ cooperatively and is able to activate CaMKII more potently if it has more *Ca*^2+^ ions bound to it [85].
- **Neurogranin (Ng)-CaM interactions**: In the absence of *Ca*^2+^, CaM is bound to scaffolding protein neurogranin (Ng), which dramatically reduces its affinity for *Ca*^2+^ (Figure 1B). On the other hand, *Ca*^2+^ decreases binding affinity of CaM for Ng [40]. Thus, CaM activation and therefore CaMKII activation depend on the competitive effects of *Ca*^2+^ and Ng.
- **The role of structure in CaMKII activation**: Further complexity for CaMKII activation is built into the structure of the molecule itself. CaMKII is a dodecamer arranged in 2 stacked hexomeric rings [9, 20]. The monomers of CaMKII (mCaMKII) consist of the kinase domain, CaM-binding domain, and phosphorylation sites T286(287), and T305(306) [81, 35, 55, 36, 72]. When the CaM-binding domain is unoccupied, and the T286(287) site is unphosphorylated, the monomer is in a conformation such that the kinase is inactive (Figure 1C, top). When CaM is bound to a CaMKII monomer, the latter undergoes a conformational change, such that the kinase is now active. Active CaMKII can phosphorylate other *Ca*^2+^*/CaM* -bound CaMKII monomers resulting in CaMKII autophosphorylation. A CaMKII monomer can only be phosphorylated if the calmodulin bound to it has at least one calcium ion, and once phosphorylated, it remains in the active state even after unbinding from CaM (Figure 1C middle). In this manner, a brief *Ca*^2+^ influx initiates a prolonged CaMKII activation [59, 60, 34, 42, 55]. Within the holoenzyme each of the monomers can phosphorylate neighbors, as long as the former is active and the latter has CaM bound to it (Figure 1)D [35, 80]. The phosphorylation rate depends on how many *Ca*^2+^ ions are bound to the CaM bound to the substrate [74].
- **Phosphatases are important**: Various types of protein phosphatases (PP1, PP2A, PP2B, PP2C) can dephosphorylate CaMKII in the brain and, if CaM is not bound to the latter, bring it back to its inactive state [72, 21]. In the PSD however, mainly protein phosphatase 1 (PP1) has been shown to be responsible for CaMKII dephosphorylation [21, 90] (Figure 1C bottom).

Here we sought to examine how the competition between *Ca*^2+^-mediated CaM activation and Ng scaffolding of CaM affects the response of CaMKII to calcium signals. To do so, we developed a computational model that accounts for both CaMKII monomeric activation and the holoenzyme kinetics in an agent-based framework. This model is built on a previously published model of CaMKII monomers activation by *Ca*^2+^*/CaM* [74] and includes the role of the scaffolding molecule Ng and the protein phosphatase PP1. Finally, we build a model of a dodecameric holoenzyme, and study the effects of multiple *Ca*^2+^ spikes on CaM-CaMKII-PP1 system in the presence and absence of Ng. Using this model, we investigated the dynamics of CaMKII phosphorylation as a function of *Ca*^2+^-influx dynamics and interactions with Ng. Results from our model show that CaMKII behaves as a leaky integrator and more importantly, the scaffold molecule, Ng, tunes the behavior of the leaky integrator.

**Figure 1:**
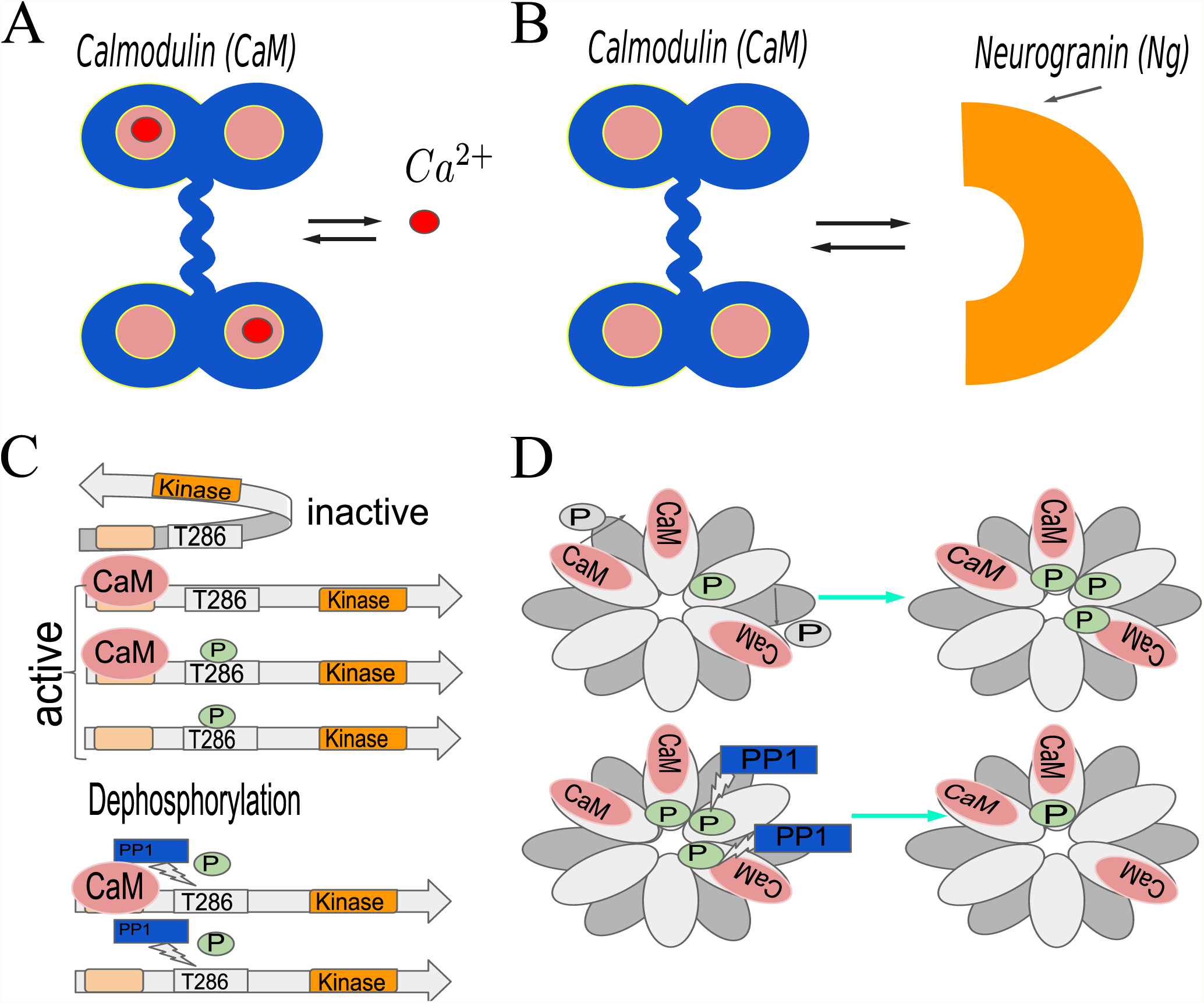
Schematic representation of the interactions between calmodulin (CaM), calcium, CaMKII, and neurogranin (Ng). (A) Calmodulin has 4 calcium binding sites; it binds *Ca*^2+^ cooperatively and its ability to activate CaMKII depends on how many of these sites are occupied. (B) Neurogranin (Ng) is a scaffolding molecule; upon binding calmodulin it dramatically reduces the latter’s affinity for calcium. In this work, we assume that CaM cannot bind calcium and Ng simultaneously. (C) The default conformation of CaMKII monomers is inactive (top); they can be activated by CaM binding. Once bound to CaM, the monomers can be phosphorylated by another active CaMKII protein. In this case, the CaMKII monomer will remain active even after losing CaM. Protein phosphatase 1 (PP1) dephosphorylates CaMKII (bottom). (D) CaMKII holoenzyme is a dodecamer, which consists of 2 stacked hexomeric rings of CaMKII monomers. Within the ring, a given CaMKII monomer can be phosphorylated by its neighbor provided that they are both in the active conformation. This is commonly referred to as autophosphorylation.

## 2 Methods

We constructed the models at different scales to characterize CaMKII phosphorylation at increasing levels of complexity. First, we added the scaffolding molecule Ng to the model from *Pepke et al* [74], to investigate the effect of Ng on CaMKII phosphorylation dynamics. Second, we added PP1 to this CaMKII monomer model to simulate the phosphorylation-dephosphorylation cycle and characterize the effects of Ng on this system. Finally, we built a model of a dodecameric holoenzyme and looked at the response of the holoenzyme-phosphatase system to calcium signals and how it is affected by Ng. An important assumption of all the models developed in this study is that only one of the phosphorylation sites (T286/7) of CaMKII is considered throughout. The phosphorylation of T305/6 site is a slower reaction that is known to inhibit CaM binding to T286/7-unphosphorylated CaMKII subunit [23], and is omitted from our simulations.

### 2.1 Model description

#### 2.1.1 Model of CaMKII monomers

We begin with the model of CaMKII monomers (mCaMKII) described by Pepke et al. [74]. This model includes *Ca*^2+^ binding rates to CaM, CaM binding rates to mCaMKII, and phosphorylation rates of active CaMKII monomers by one another, all depending on how many *Ca*^2+^ ions are bound to the CaM molecules involved. We incorporate Ng binding to CaM with the rate constants from [79], and assume that the binding of CaM to *Ca*^2+^ and Ng is mutually exclusive (Table 1). In addition to the reactions in [74], we included CaM unbinding and binding to phosphorylated CaMKII, albeit with slower kinetics [58]. This reaction is important for the timescales of our interest (on the order of minutes) and we adapt these reaction rates from [58]. Finally, CaMKII dephosphorylation by PP1 is modeled as Michaelis-Menten kinetics, with the rate constants from [16] (Table 1).

**Table 1:** Reaction rates for the model. All the numbers with the exception of “CaMKII binding to CaM-CaMKII” are used in both monomers and holoenzyme models. The difference between the 2 models are the initial conditions: we either start the simulations with only monomers or only holoenzyme, and do not allow the disintegration of the latter. * only present in the model for monomers

| Parameter | Value | Reference |
| --- | --- | --- |
| $Ca^{2+}$ binding to CaM | | |
| $k_{on}^{1C}$ | $4 \mu M^{-1}s^{-1}$ | [74, 53, 50, 85] |
| $k_{off}^{1C}$ | $40.24 s^{-1}$ | [75, 57, 33] |
| $k_{on}^{2C}$ | $10 \mu M^{-1}s^{-1}$ | [74, 53, 50, 85] |
| $k_{off}^{2C}$ | $9.3 s^{-1}$ | [33] |
| $k_{on}^{1N}$ | $100 \mu M^{-1}s^{-1}$ | [74, 53, 50, 85] |
| $k_{off}^{1N}$ | $2660 s^{-1}$ | [75, 33, 73, 17] |
| $k_{on}^{2N}$ | $150 \mu M^{-1}s^{-1}$ | [74, 53, 50, 85] |
| $k_{off}^{2N}$ | $990 s^{-1}$ | [75, 57, 33, 73, 17] |
| CaM binding to unphosphorylated CaMKII |  |  |
| $k_{on}^{CaM0}$ | $3.8 \cdot 10^{-3} \mu M^{-1}s^{-1}$ | [74, 58] |
| $k_{off}^{CaM0}$ | $6.56 s^{-1}$ | [74, 58] |
| $k_{on}^{CaM1C}$ | $59 \cdot 10^{-3} \mu M^{-1}s^{-1}$ | [74, 58] |
| $k_{off}^{CaM1C}$ | $6.72 s^{-1}$ | [74, 58] |
| $k_{on}^{CaM2C}$ | $0.92 \mu M^{-1}s^{-1}$ | [74, 58] |
| $k_{off}^{CaM2C}$ | $6.35 s^{-1}$ | [74, 58] |
| $k_{on}^{CaM1C1N}$ | $0.33 \mu M^{-1}s^{-1}$ | [74, 58] |
| $k_{off}^{CaM1C1N}$ | $5.68 s^{-1}$ | [74, 58] |
| $k_{on}^{CaM2C1N}$ | $5.2 \mu M^{-1}s^{-1}$ | [74, 58] |
| $k_{off}^{CaM2C1N}$ | $5.25 s^{-1}$ | [74, 58] |
| $k_{on}^{CaM1N}$ | $22 \cdot 10^{-3} \mu M^{-1}s^{-1}$ | [74, 58] |
| $k_{off}^{CaM1N}$ | $5.75 s^{-1}$ | [74, 58] |
| $k_{on}^{CaM2N}$ | $0.1 \mu M^{-1}s^{-1}$ | [74, 58] |
| $k_{off}^{CaM2N}$ | $1.68 s^{-1}$ | [74, 58] |
| $k_{on}^{CaM1C2N}$ | $1.9 \mu M^{-1}s^{-1}$ | [74, 58] |
| $k_{off}^{CaM1C2N}$ | $2.09 s^{-1}$ | [74] |
| $k_{on}^{CaM4}$ | $30 \mu M^{-1}s^{-1}$ | [74, 58] |
| $k_{off}^{CaM4}$ | $1.95 s^{-1}$ | [74, 58] |
| $Ca^{2+}$ binding to CaM-CaMKII | | |
| $k_{on}^{K1C}$ | $44 \mu M^{-1}s^{-1}$ | [74, 33] |
| $k_{off}^{K1C}$ | $29.04 s^{-1}$ | [74, 33] |
| $k_{on}^{K2C}$ | $44 \mu M^{-1}s^{-1}$ | [74, 33] |
| $k_{off}^{K2C}$ | $2.42 s^{-1}$ | [74, 33] |
| $k_{on}^{K1N}$ | $75 \mu M^{-1}s^{-1}$ | [74, 33] |
| $k_{off}^{K1N}$ | $301.5 s^{-1}$ | [74, 33] |
| $k_{on}^{K2N}$ | $76 \mu M^{-1}s^{-1}$ | [74, 33] |
| $k_{off}^{K2N}$ | $32.68 s^{-1}$ | [74, 33] |
| CaMKII binding to CaM-CaMKII * |  |  |
| $k_{on}^{CaMKII}$ | $50 \mu M^{-1}s^{-1}$ | [74, 22] |
| $k_{off}^{CaMKII}$ | $60 s^{-1}$ | [74, 35, 15] |
| CaMKII phosphorylation |  |  |
| $k_p^{CaM0}$ | $0 s^{-1}$ | [74, 85] |
| $k_p^{CaM1C}$ | $0.032 s^{-1}$ | [74, 85] |
| $k_p^{CaM2C}$ | $0.064 s^{-1}$ | [74, 85] |
| $k_p^{CaM1C1N}$ | $0.094 s^{-1}$ | [74, 85] |
| $k_p^{CaM2C1N}$ | $0.124 s^{-1}$ | [74, 85] |
| $k_p^{CaM1N}$ | $0.061 s^{-1}$ | [74, 85] |
| $k_p^{CaM2N}$ | $0.12 s^{-1}$ | [74, 85] |
| $k_p^{CaM1C2N}$ | $0.154 s^{-1}$ | [74, 85] |
| $k_p^{CaM4}$ | $0.96 s^{-1}$ | [74, 85] |
| CaM binding to phosphorylated CaMKII |  |  |
| $k_{p,on}^{CaM0}$ | $1.27 \cdot 10^{-3} \mu M^{-1}s^{-1}$ | [74, 58] |
| $k_{p,on}^{CaM1C}$ | $19.7 \mu M^{-1}s^{-1}$ | [74, 58] |
| $k_{p,on}^{CaM2C}$ | $0.3 \mu M^{-1}s^{-1}$ | [74, 58] |
| $k_{p,on}^{CaM1C1N}$ | $1.1 \mu M^{-1}s^{-1}$ | [74, 58] |
| $k_{p,on}^{CaM2C1N}$ | $1.73 \mu M^{-1}s^{-1}$ | [74, 58] |
| $k_{p,on}^{CaM1N}$ | $7.3 \mu M^{-1}s^{-1}$ | [74, 58] |
| $k_{p,on}^{CaM2N}$ | $0.03 \mu M^{-1}s^{-1}$ | [74, 58] |
| $k_{p,on}^{CaM1C2N}$ | $0.63 \mu M^{-1}s^{-1}$ | [74, 58] |
| $k_{p,on}^{CaM4}$ | $10 \mu M^{-1}s^{-1}$ | [74, 58] |
| $k_{p,off}^{CaM}$ | $0.07 s^{-1}$ | [58] |
| Ng binding to CaM |  |  |
| $k_{on}^{Ng}$ | $5 \mu M^{-1}s^{-1}$ | [79] |
| $k_{off}^{Ng}$ | $1 s^{-1}$ | [79] |
| CaMKII dephosphorylation by PP1 (Michaelis-Menten constants) |  |  |
| $k_{cat}$ | $0.41 s^{-1}$ | [16] |
| $K_m$ | $11 \mu M$ | [16] |

#### 2.1.2 Model of CaMKII holoenzyme

##### Assumptions specific to the holoenzyme model

It has been shown that while the kinase domains of individual subunits are attached to the rigid hub domain with a highly flexible linker domain, ∼ 20% of subunits form dimers, and ∼ 3% of them are in a compact conformation, both of which render CaM-binding-domain inaccessible [62]. Here, we do not include such detailed interactions between linker domains and flexibility of the individual subunits within the holoenzyme. Rather, we assume that the kinase domains are positioned rigidly within the 2 hexameric rings, such that each subunit is in a position to phosphorylate only one of its neighbors as depicted in Figure 1D.

We further assume that the CaMKII subunits within the holoenzyme have the same binding rates to different species of *Ca*^2+^*/CaM* as the CaMKII monomers, and once activated their phosphorylation rate is the same as that of the corresponding monomers. Based on these assumptions, the model for the CaMKII holoenzyme does not include the reaction for 2 CaMKII monomers binding to one another. Rather, once the appropriate neighbor of an active CaMKII subunit binds *Ca*^2+^*/CaM*, the phosphorylation rule is invoked with the corresponding reaction rate (Table 1).

##### Model development

To model the dynamics of CaMKII holoenzyme, we changed the types of molecules that we start the simulation with. Instead of CaMKII monomers, we initiated the simulation with CaMKII holoenzymes, and defined the phosphorylation rules for individual subunits such that a given subunit can only phosphorylate one of its neighbors (see model assumptions). The rules are the same as those of the monomers, with the exception of 2 CaMKII monomers binding each other to phosphorylate one another, since they are already bound within the holoenzyme. In this case, the reaction rules apply to the CaMKII subunits within the holoenzyme rather than CaMKII monomers.

This model does not contain any CaMKII monomers outside of the holoenzyme. We calculated the total number of the holoenzymes to keep the concentration of the [*mCaMKII*] = 80 μ*M* consistent with our model of the monomers. For our simulated PSD volume *V* = 0.0156 μm^3^ this concentration corresponds to 80 μM = 80 · (6.022 · 10^23^ · 0.0156 · 10^−15^) · 10^−6^ monomers which constitutes 752*/*12 ≈ 63 holoenzymes.

### 2.2 Rule-Based modeling and BioNetGen

Since each CaMKII monomer can bind a calmodulin molecule, which can adopt 9 distinct states (assuming that the 2 *Ca*^2+^ binding sites on each of the lobes are indistinguishable), and be phosphorylated or unphosphorylated, there are more than 18^12^ states the dodecameric holoenzyme could adopt. To avoid a combinatorial explosion of states we built our model in a rule-based modeling language BioNetGen [37], which is briefly described here.

A powerful idea at the heart of rule-based modeling is colloquially referred as “don’t care, don’t write” [28, 13]. Here, for a given reaction (or rule), we only specify the states of the reactants that are relevant for the said reaction, and leave the rest unspecified. For example, a specific CaMKII subunit will bind a given species of *Ca*^2+^*/CaM* with the same rate regardless of the conformation of its neighbors. Thus, a rule to bind a CaM molecule that has 2 *Ca*^2+^ ions bound to its C-terminus would look like:

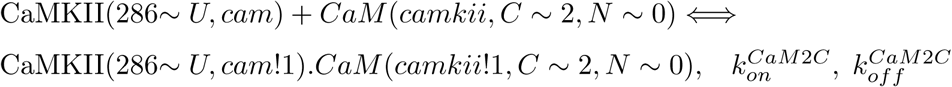

This rule indicates that regardless of the presence or conformations of a non-phosphorylated (indicated by 286 ∼ *U*) CaMKII subunits’ neighbors it binds CaM, with 2 *Ca*^2+^ ions at its C lobe, reversibly with the reaction rates 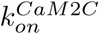 and 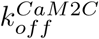 (Table 1). This dramatically reduces the number of reactions that need to be written. In the case of our CaMKII model, we only end up with ∼ 40 reactions despite the huge number of possible states.

In BioNetGen, once the reactions have been specified, if there is no danger of running into combinatorial explosion (the numbers of possible states and reactions are manageable), a full network of reactions can be generated, and the simulations can be run with ordinary differential equations (ODEs) [13]. If the number of states of a given enzyme is too large however, a stochastic agent-based approach is adopted, in which case only the reactions between existing discrete molecules that occur during the simulation need to be tracked by the program (network free simulation [89, 92]). In this case, the important quantity is the number of different states present which can be much smaller than the number of possible states. In this work, we conduct the simulations for the monomer model with ODEs, and for the holoenzyme model with the network free method.

## 3 Results

### 3.1 mCaMKII phosphorylation dose response curve depends on Ng non-linearly

Since Ng is a scaffolding molecule and its presence is expected to decrease the [*CaM*]_*free*_ : [*Ca*^2+^]_*free*_ ratio we first investigated how it affects the phosphorylation of CaMKII at different [*CaM*] while maintaining [*Ca*^2+^] constant. Following Pepke *et al.* [74], we set [*CaMKII*] = 80 μM and calculated the dose response to calmodulin at [*Ca*^2+^] = 10 μM with and without Ng. When present, the concentration of Ng was set to 20 μM [101].

Since CaMKII phosphorylation depends on CaM, and the activity of the latter depends on how many *Ca*^2+^ ions it binds, we first look at representations of different *Ca*^2+^*/CaM* species depending on [CaM]. Figure 2 depicts the maximum amounts of each individual *Ca*^2+^*/CaM* species relative to total CaM available for a range of [CaM]s. We see that without Ng, we consistently have more 1*Ca*^2+^*/CaM* than in the presence of Ng (Figure 2A, and B top). This is easily understandable: Ng limits the availability of free calmodulin, and without it we have more CaM available to bind calcium. On the other hand, when we look at CaM species that have multiple calcium ions bound to them, we see a crossover of the 2 curves as the total [CaM] increases. This is particularly striking for 3*Ca*^2+^*/CaM* and 4*Ca*^2+^*/CaM* (Figures 2C, B and E bottom). For these species, limiting the available free CaM can be beneficial since this increases the ratio [*Ca*^2+^] : [*CaM*]_*free*_, resulting in higher number of calmodulin proteins bound to multiple calcium ions. Figure 2E shows the dose response in the physiological range of CaM concentration. This might seem like a small effect, but it is amplified by the fact that these multi-calcium-bound calmodulin species have a higher CaMKII binding rate, and render the bound CaMKII more susceptible to phosphorylation (Table 1).

**Figure 2:**
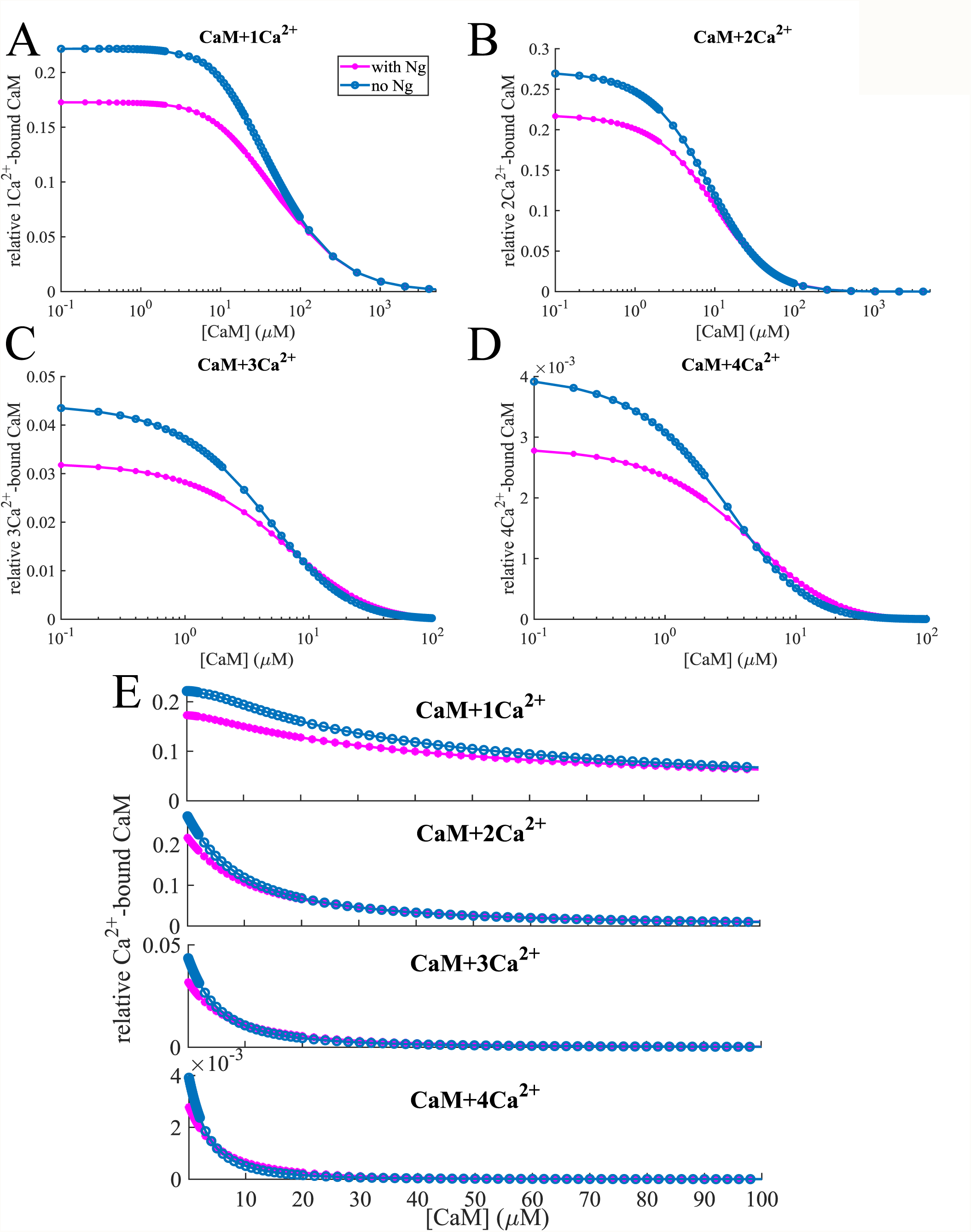
Relative concentration of (A) 1-, (B) 2-, (C) 3- and (D) 4-calcium-bound calmodulin vs. total [CaM]. For most of the calcium-bound forms, the relative concentration is lower in the presence of neurogranin. However, in the case of 3- and 4-calcium-bound calmodulin the situation is reversed for certain [CaM]: Ng competes with *Ca*^2+^ for free calmodulin, which allows a regime where the concentration of these “multi-calcium-bound” forms is higher. As the concentration of calmodulin is further increased, the [*Ca*^2+^] : [*CaM*] ratio decreases and these forms die out, thus at high [CaM] the effects of Ng are diminished. (E) shows the same effect for physiologically relevant [CaM].

When we look at phosphorylated CaMKII bound to different species of calmodulin, we see that Ng has very different effects on these different species (Figure 3). Phosphorylation by 1- and 2-*Ca*^2+^*/CaM* in the presence of Ng is lower than that in the absence of Ng for [*CaM*] ≈ 1 − 100 μM. (Figures 3A, B, and E top 2). As [CaM] increases, the presence of Ng becomes less relevant and the 2 curves converge. The situation is quite different for 3- and more importantly 4-*Ca*^2+^*/CaM* species (Figures 3C, D and E bottom 2). At slightly higher [CaM], the competition with Ng for free CaM plays an important role: a higher proportion of the CaM molecules have 4 calcium ions bound to them in the presence of Ng than in the absence as discussed above, and these CaM molecules give a much higher phosphorylation rate to the CaMKII monomers that they bind. However, these species only exist at relatively lower [CaM]s – as [CaM] increases, the ratio [*Ca*^2+^] : [*CaM*]_*free*_ decreases, the effect of Ng becomes insubstantial since the 1- and 2-*Ca*^2+^*/CaM* species are responsible for CaMKII phosphorylation at these concentrations. The dose response curves in the physiologically relevant range of [CaM] (∼ 10 − 50 μM, [12, 44, 96], Figure 3E) show that the influence of Ng on CaMKII phosphorylation is most prominent in this range.

**Figure 3:**
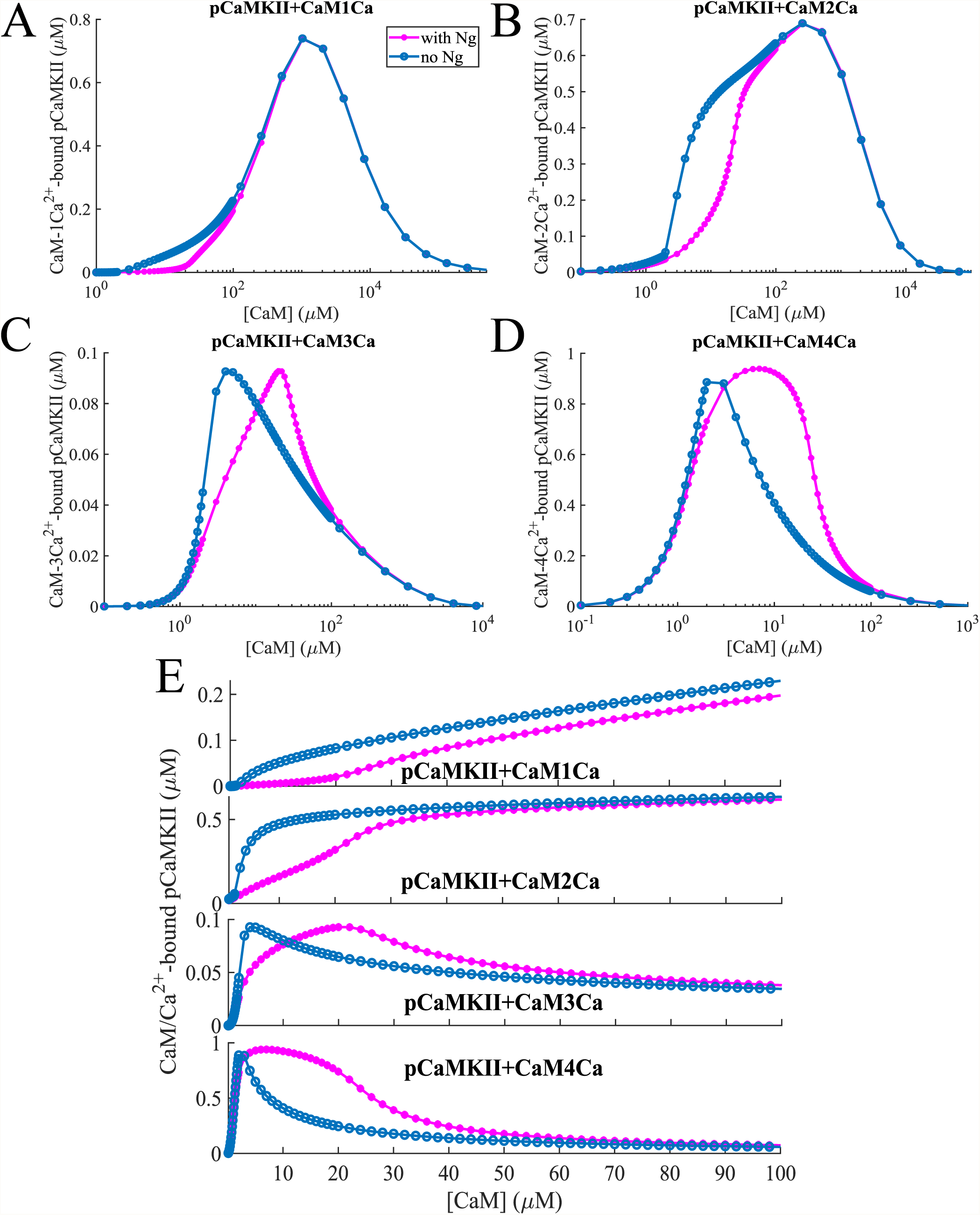
Dose response of phosphorylated CaMKII molecules bound to different states of CaM-*Ca*^2+^ as shown on Figure 2. At low [CaM], the “multi-calcium-bound” forms of calmodulin play an important role. This is amplified by the fact that these forms when bound to CaMKII give it higher phosphorylation rates (Table 1).

The dose response of total CaMKII phosphorylation to CaM is depicted in Figure 4A. Here, the first local maximum is caused by phosphorylation of the CaMKII monomers bound to 4 − *Ca*^2+^*/CaM*, and since there is more of these species in the presence of Ng, this peak is more prominent when Ng is present in the system. The second peak is caused by phosphorylation of the CaMKII monomers bound to 1- and 2- *Ca*^2+^*/CaM*, which play an important role at higher [CaM] where the role of Ng is less important, and so the 2 curves increasingly overlap as [CaM] is increased. The implications of this result can be understood by comparing Figures (4B and C), which show the dynamics of CaMKII phosphorylation with and without Ng at 2 distinct [CaM]s, an average (30 μM) and an upper bound to physiological (100 μM). The effect of Ng is reversed in the case of these 2 CaM concentrations: when [*CaM*] = 30 μM, Ng increases CaMKII phosphorylation level, and when [*CaM*] = 100 μM, Ng decreases the overall phosphorylation of CaMKII. Thus, Ng affects the dose response of CaMKII phosphorylation to CaM in a non-linear manner and these effects can be clearly understood by examining the effect of this scaffolding molecule on *Ca*^2+^*/CaM* species and their ability to activate CaMKII.

**Figure 4:**
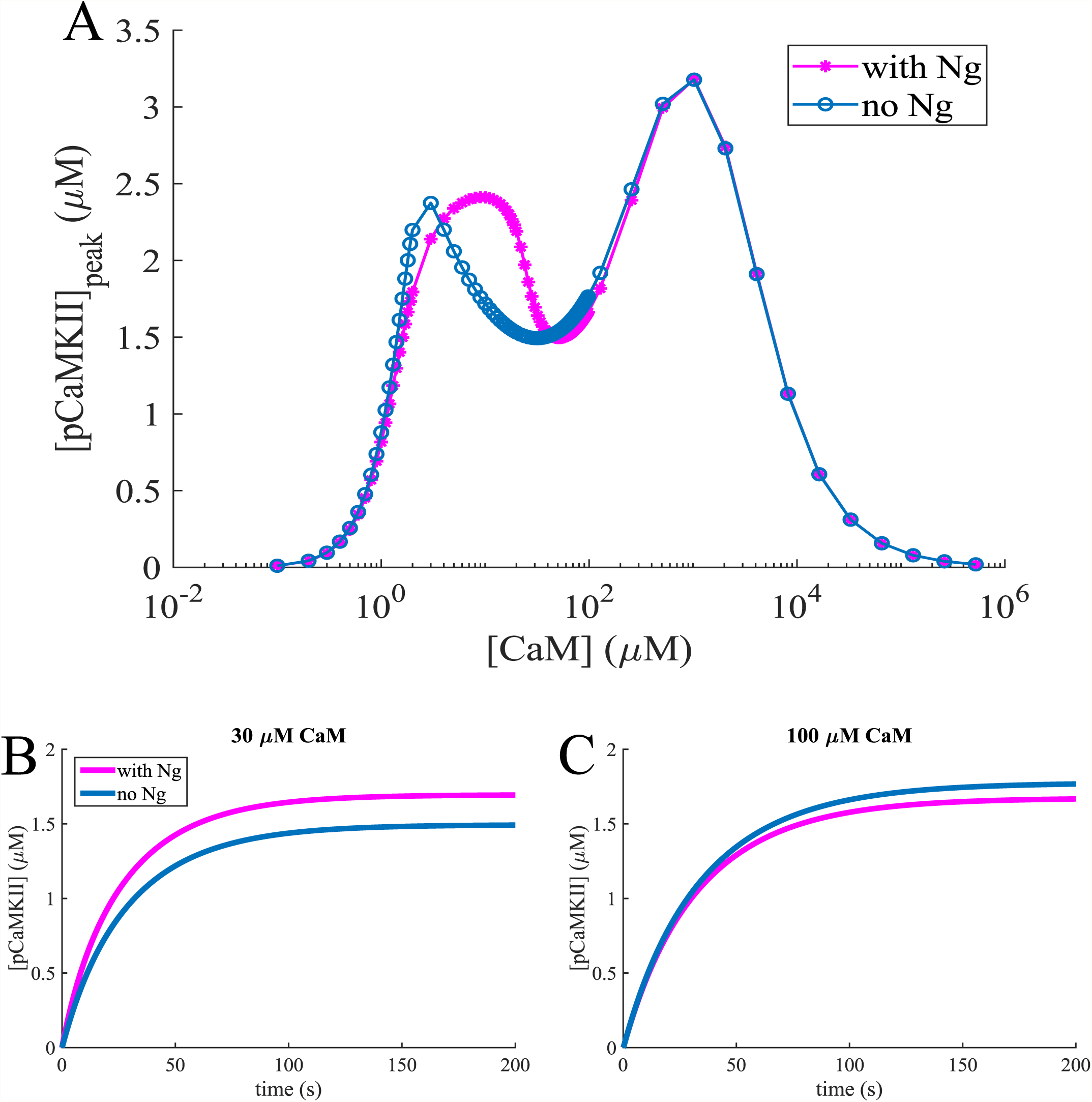
CaMKII phosphorylation dependence on [CaM] and Ng. (A) Steady state phosphorylated CaMKII concentration as a function of total CaM concentration in the presence of 10 μM *Ca*^2+^ with (pink lines) and without (blue lines) Ng. (B) and (C) Dynamics of CaMKII monomers (80 μM) autophosphorylation and dephosphorylation by PP1, in the presence of constant [*Ca*^2+^] = 10 μM with and without Ng, with [*CaM*] = 30 μM and [*CaM*] = 100 μM respectively.

### 3.2 Ng modulates mCaMKII phosphorylation in response to a> *Ca*^2+^ spike

In dendritic spines, the steady state calcium concentration is ∼ 100*nM* [82]. Any *Ca*^2+^ influx is rapidly buffered in the spine, so that [*Ca*^2+^]_*free*_ falls back to near steady state levels within ∼ 100 ms [31]. To better mimic the experiments, we adjust the total amount of *Ca*^2+^ in the spine so that the free *Ca*^2+^ ions reach ∼ 10 μM at the peak of the signal and fall back to near steady state levels within ∼ 100 ms [100, 26, 76, 5, 31, 63]. Note that this is not representative of the actual amount of calcium ions entering the spine, since our model does not include any of the calcium binding molecules (with the exception of calmodulin), and only simulates calcium buffering mathematically.

Before simulating the calcium spike, we allow the system to equilibrate. There is some basal level of CaMKII phosphorylation even at the low calcium concentrations (∼ 100 nM) observed in the spine in equilibrium. To achieve a given [*Ca*^2+^]_*free*_ in different conditions, the concentration of total calcium varies depending on the conditions simulated, and so does the basal CaMKII phosphorylation level. To enable direct comparison, we look at the difference of the peak and basal phosphorylation for each condition, as well as the Area Under the Curve (AUC), peak time, and the decay time of phosphorylation levels.

We first investigated the dependence of maximum CaMKII phosphorylation on the amplitude of the [*Ca*^2+^]_*free*_ pulse. In Figure 5, the horizontal axis represents the “measured” [*Ca*^2+^]_*free*_ and the vertical axis shows the level of maximal CaMKII phosphorylation. As we can see in this case, for any [*Ca*^2+^]_*free*_ the response is higher in the absence of Ng, indicating that even for very small [*Ca*^2+^] spikes, the availability of CaM is the limiting factor for CaMKII phosphorylation. This is further confirmed by the fact that as the size of the *Ca*^2+^ spike increases, the maximum CaMKII phosphorylation levels in the presence and absence of Ng diverge from each other as they reach saturation.

**Figure 5:**
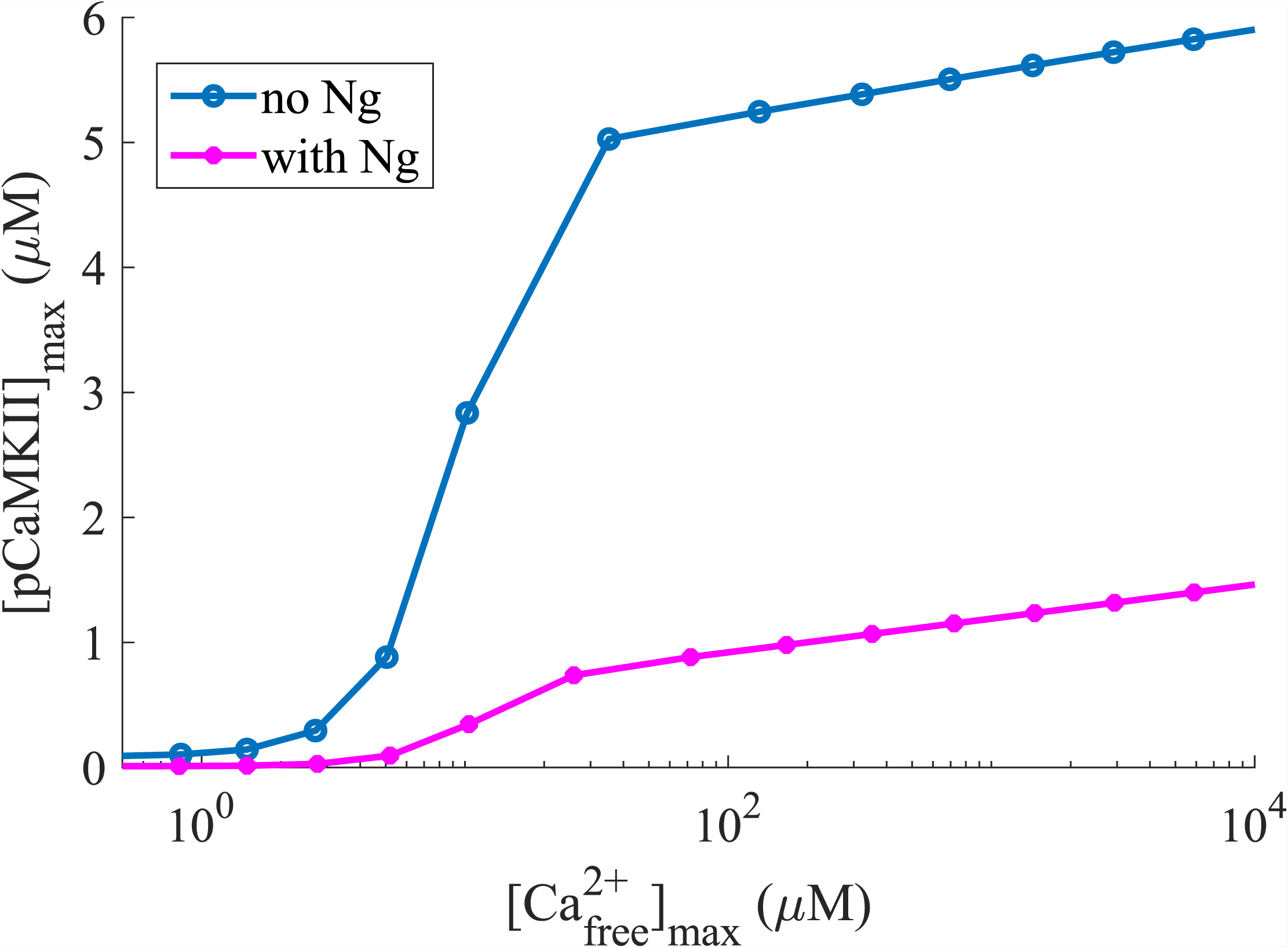
Maximum phosphorylated CaMKII concentration for different concentrations of *Ca*^2+^ spikes with [CaM] = 30 μM. The horizontal axis represents the free calcium available at the peak of the spike.

The dynamics of CaMKII phosphorylation is shown on Figure 6A for a range of physiological CaM concentrations including the upper limit of 100 μM. As we can see, even in this extreme case Ng decreases CaMKII phosphorylation, indicating the [CaM] is the limiting factor. Interestingly, the phosphorylation dynamics with Ng and at 50 μM [CaM] is identical to that without Ng at 30 μM [CaM].

**Figure 6:**
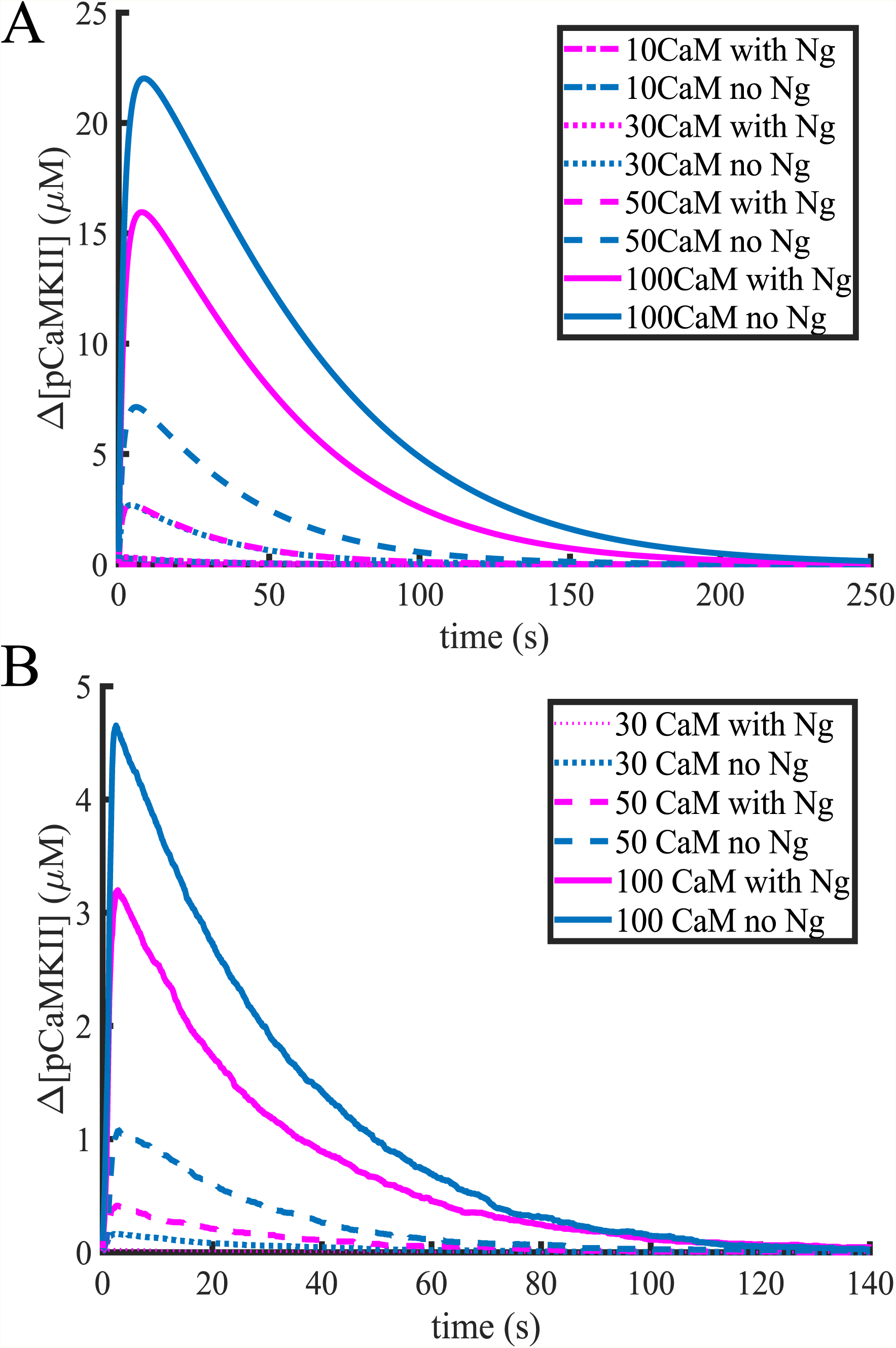
The rise in CaMKII phosphorylation level as a response to a [*Ca*^2+^]_*free*_ = 10 μM spike with monomers (A) and holoenzymes (B) for different conditions. The response of the holoenzyme at [CaM] = 10 μM was not significant and is not shown here.

Further comparing the total amount of CaMKII phosphorylation over time (Area Under the Curve or AUC) as well as the relative peak phosphorylation levels (Figures 7A, top and B), we see that the presence of Ng makes a striking difference – Ng dramatically impairs phosphorylation of CaMKII. However, at extreme high [CaM] (100 μM), the system is robust to the presence of Ng although the latter still modulates the level of phosphorylation. The presence of Ng does not appear to have a dramatic effect on the peak time or phosphorylation lifetime, which we define as the time it takes to attain 10% of the peak phosphorylation level (Figures 7E, and G). This is expected; once phosphorylated, the CaMKII molecules can be dephosphorylated by PP1, which works independently of Ng (and in our model, of any other molecule).

**Figure 7:**
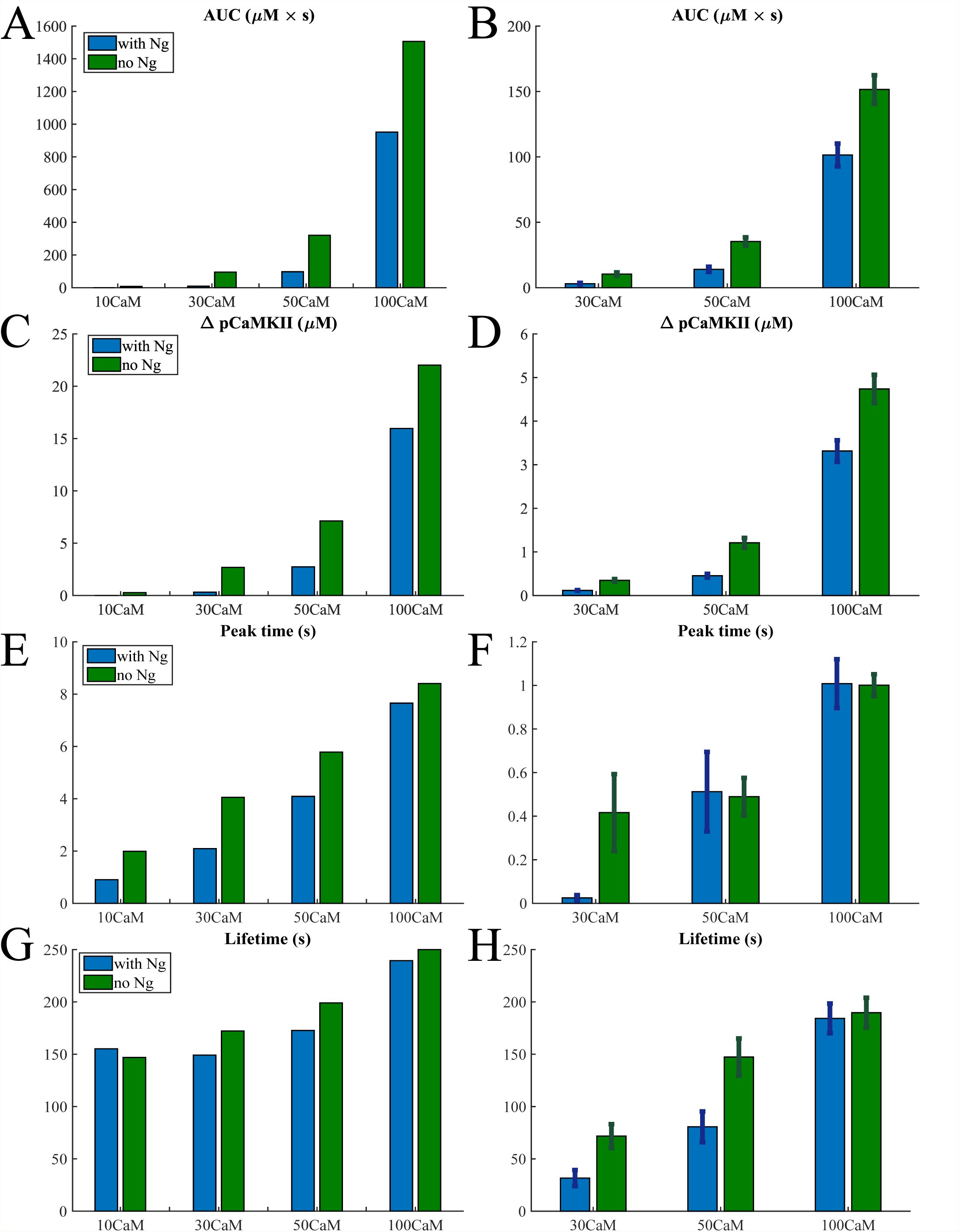
Bar graphs in different conditions for the monomers and holoenzymes respectively of Area Under the Curve (AUC) (A) and (B), the maximum increase in phosphorylated CaMKII concentration (C) and (D), the time of the phosphorylation level reaching its peak (E) and (F), and the lifetime defined in the main text (G) and (H) in response to a 10 μM *Ca*^2+^ pulse.

### 3.3 CaMKII holoenzyme phosphorylation is robust against fluctuations of [*Ca*^2+^]_*free*_

We next investigated the role of Ng in governing the phosphorylation dynamics of the CaMKII holoenzyme using a stochastic agent-based approach. Figure 6B shows the average CaMKII phosphorylation in response to a ∼ 10 μM single *Ca*^2+^ spike for [*CaM*] = 30 μ*M,* 50 μ*M*, and 100 μ*M*, respectively. We also conducted simulations at [*CaM*] = 10 μ*M* ; however at this CaM concentration, CaMKII did not react to 10 μM *Ca*^2+^ pulses. The results shown are an average of ∼ 30 runs of the stochastic model. In the simulations with [*CaM*] = 30 μM, the mean peak and standard deviation of [*Ca*^2+^]_*free*_ were 10.7 ± 1.6 μM in the case with no Ng and 10.3 ± 1.2 μM in the case with Ng. We note that with this [CaM], CaMKII would not always react to the 10 μM free calcium spike, and sometimes there would be no detected phosphorylated CaMKII. These measurements were not taken into account in the calculations plotted on Figures 6B and 7B, D, F and H. For the model without Ng, 36 out 60 simulations yielded a change in phosphorylation levels. For the model with Ng, we had to conduct 120 simulations to get as many as 31 that resulted in noticeable CaMKII phosphorylation. From these observations, we can conclude that the probability of CaMKII phosphorylation is affected by the presence of Ng. When [*CaM*] = 50 μM, the mean peak and standard deviation of [*Ca*^2+^]_*free*_ were 9.4 ± 1.2 μM in the case with no Ng and 10.2 ± 1.3 μM in the case with Ng. Finally, in the extreme case when [*CaM*] = 100 μM the corresponding numbers were 9.2 ± 1.3 μM both with and without Ng.

By comparing the Figures 6A and B, 7A, C, E and Figures 7B, D and F, we notice that with the same amount of CaM present in the simulation, the holoenzyme reacts significantly faster to the calcium signal (see the peak time) than the monomeric CaMKII, however the phosphorylation level is also much lower. This tells us that the holoenzyme is more robust to *Ca*^2+^ fluctuations than the monomers. When the holoenzyme does react to a calcium signal, however, the phosphorylation lifetime is comparable to that of the monomers (Figures 7G and **??**H). And as in the case for monomers the presence of Ng modulates the effect of the calcium signal at physiological calmodulin concentrations.

### 3.4 A leaky integrator of [*Ca*^2+^] signals

To mimic experimental stimuli of a train of *Ca*^2+^ spikes [32, 91, 52, 19], we next investigated the response of the system to multiple calcium spikes. Figure 8 shows the average response of CaMKII phosphorylation to 10 calcium pulses at 0.5 Hz. As we can see, even with 10 *Ca*^2+^ pulses in the case of [*CaM*] = 30 μ*M*, the response is very small, particularly in the presence of Ng. In fact, even with 10 calcium pulses, only 25 out of 30 simulations resulted in a non-negligible difference of CaMKII phosphorylation levels in the presence of Ng, and only 28 out of 30 in the absence of Ng. Therefore, multiple spikes increase and even out the probability of CaMKII phosphorylation but not the overall phosphorylation level in the presence or absence of Ng.

**Figure 8:**
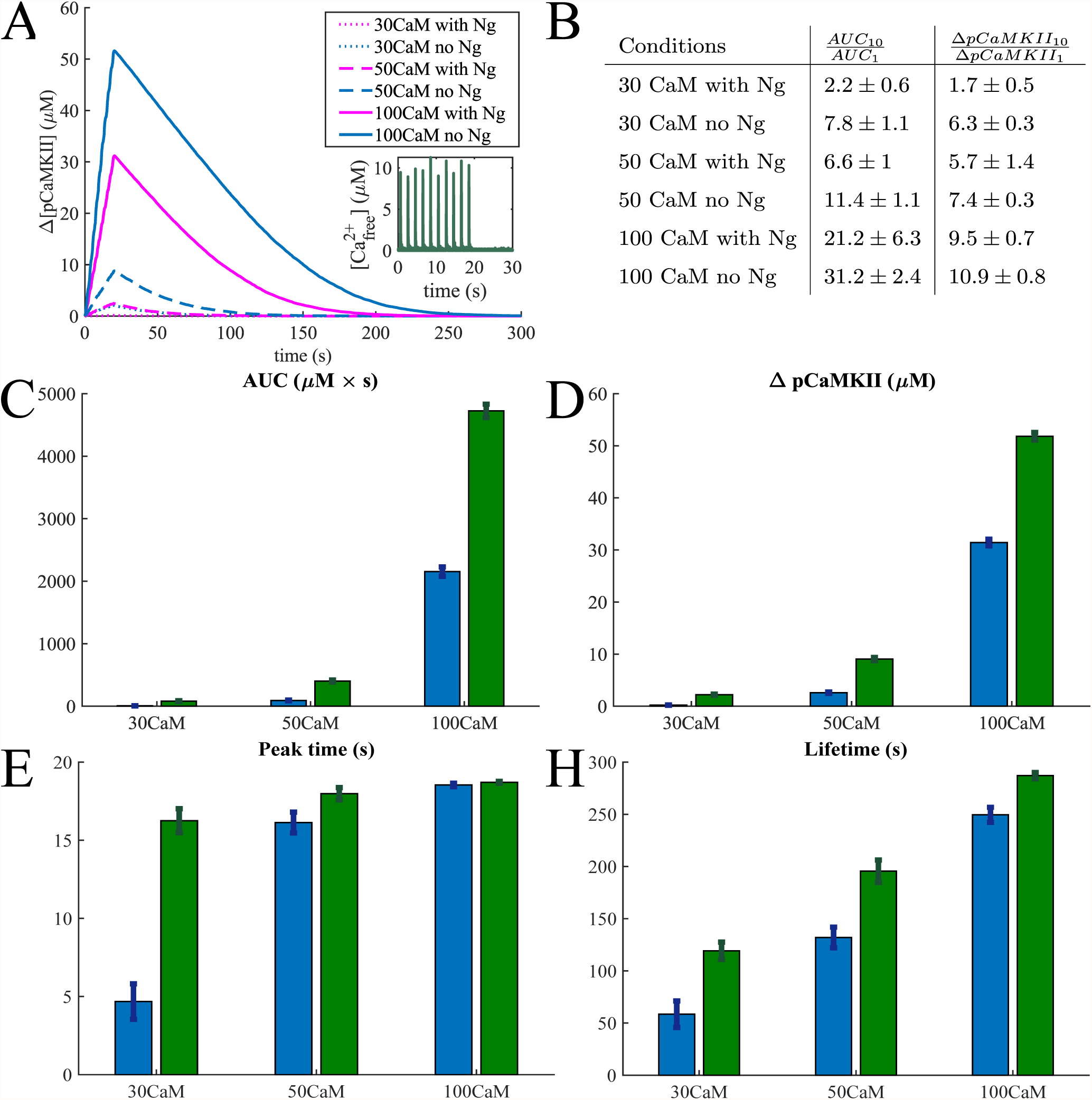
(A) An average phosphorylation of CaMKII holoenzyme in response to 10 *Ca*^2+^ pulses at 0.5 Hz averaged over 30 simulations with 63 holoenzymes and 30 μM 50 μM, and 100 μM calmodulin. (B) Table of the ratios of the Area under the curve and maximum change in phosphorylation level in response to 10 and 1 Ca pulses in different conditions. Bar graphs comparing the (C) Area Under the Curve (AUC), (D) the maximum increase in the CaMKII phosphorylation concentration, (E) the time of the phosphorylation level reaching its peak, and (F) the lifetime defined in the main text for in response these *Ca*^2+^ pulses.

As before, the presence of Ng makes less of a difference when the concentration of calmodulin is higher (100 μM), and this system is far more sensitive to calcium signals (Figures 8A, C and D). To compare the effect of multiple *Ca*^2+^ pulses versus a single pulse, we calculated the ratios of the CaMKII phosphorylation metrics, AUC and maximum change in phosphorylation for 10 to 1 calcium pulses. This ratios are shown in Figure 8B for all 6 conditions tested. As can be seen here, for average physiological conditions, a 10-fold increase in *Ca*^2+^ signal results in a more modest increase in CaMKII phosphorylation. With the exception of 100 μM (ultrahigh) CaM concentration, the ratio of the changes in phosphorylated CaMKII concentrations is consistently lower than 10, and even the ratios of AUCs get close to 10 only for [CaM] = 50 μM in the absence of Ng. This indicates a leakage in the process of the calcium signal integration to CaMKII phosphorylation.

To further investigate the leakiness of CaMKII phosphorylation, we next calculated CaMKII phosphorylation in response to 30 *Ca*^2+^ pulses (Figures 9) at average physiological [CaM]. From Figure 9A we can clearly see that the rate of increase in phosphorylation level decreases as more calcium pulses are added. This leakage of the calcium signal integration is caused by dephosphorylation of CaMKII by PP1 and has recently been observed experimentally *in vivo* [19].

**Figure 9:**
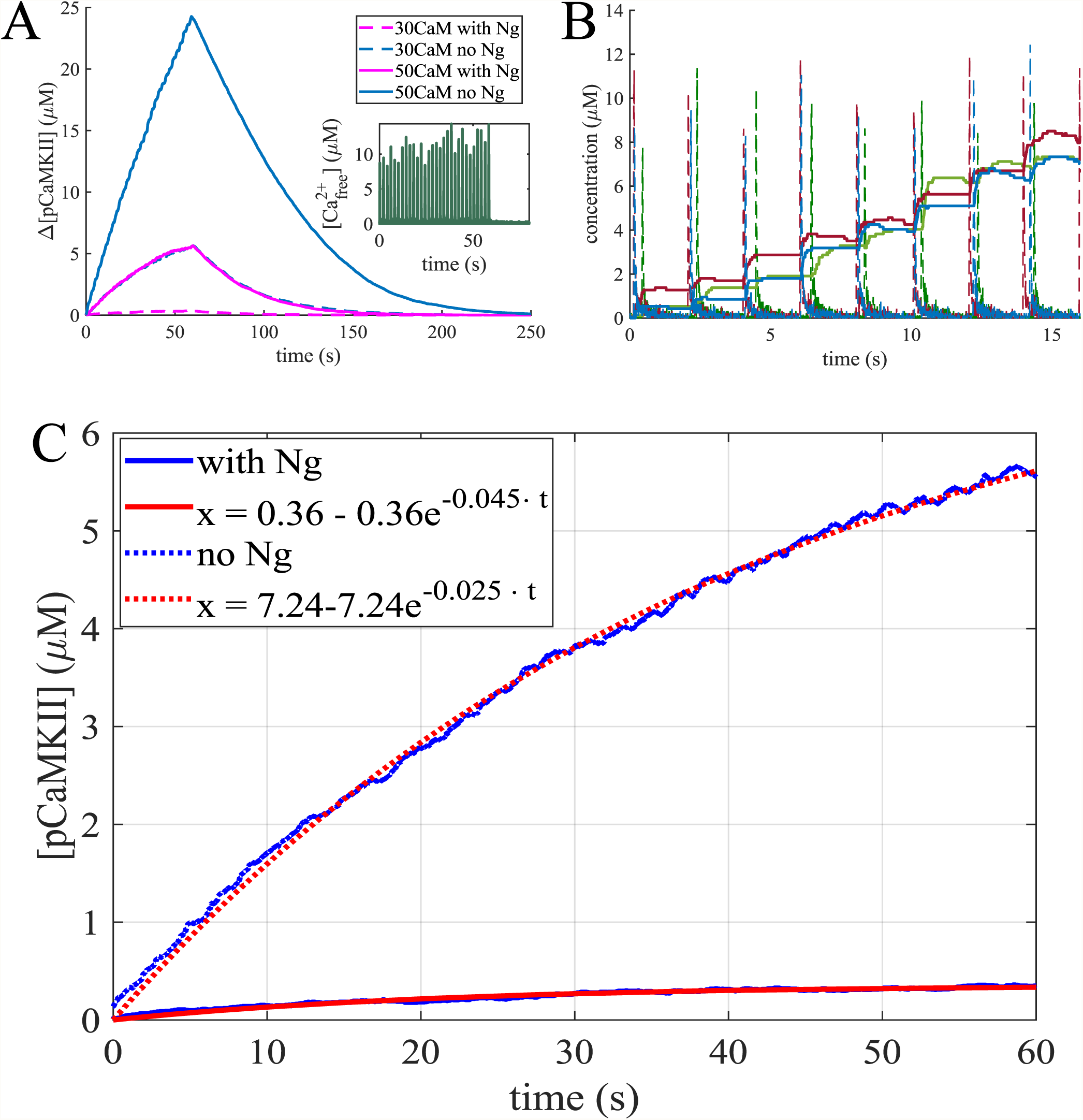
(A) An average phosphorylation of CaMKII holoenzyme in response to 30 *Ca*^2+^ pulses at 0.5 Hz averaged over 30 simulations with 63 holoenzymes and 30 μM, and 50 μM calmodulin. (B) Three individual simulations: the response of CaMKII phosphorylation level (solid lines) change to the calcium spikes (dashed lines). (C) Fitting the leaky integrator. The average response to 30 *Ca*^2+^ pulses was fitted to a curve of the form *x* = *k* − *k* · *e*^−*a·t*^ with and without Ng and [*CaM*] = 30 μM, with the curve fitting tool from MATLAB. The parameters obtained are: *k* = 7.24, *a* = 0.025 with an *R*^2^ = 0.9979 without Ng and *k* = 0.36, *a* = 0.045 with an *R*^2^ = 0.9683 with Ng. Thus Ng not only changes the leaking rate of the integrator, but significantly lowers the maximum possible phosphorylation level.

Since these simulations are conducted stochastically, the timing of *Ca*^2+^ signals is also stochastic, and not exactly synchronized across the multiple runs. Figure 9B shows CaMKII phosphorylation in response to a train of 0.5 Hz *Ca*^2+^ pulses for 3 individual runs. We see that the phosphorylation level rises in a stepwise manner in response to each calcium spike.

To better characterize CaMKII as a leaky integrator and the effect of Ng, we fit the CaMKII phosphorylation curves at a physiological [*CaM*] = 30 μM to the leaky integrator equation *x* = *k* − *k* · *e*^−*a·t*^ where k is the maximum possible change of phosphorylation level and a is the overall phosphorylation rate (Figure 7C). The curve fitting tool from *MATLAB* was used for this purpose. The response with Ng was characterized by the equation 0.36 − 0.36 · *e*^−0.045*·t*^ with an *R*^2^ = 0.9683 and the response without Ng by 7.24 − 7.24 · *e*^−0.025*·t*^ with an *R*^2^ = 0.9979. This result demonstrates that Ng not only regulates the leak rate of the integrator, but also severely reduces the maximum possible phosphorylation level. The results on Figures 8 and 9 predict that our model behaves as a leaky integrator of calcium signals, and that the presence of Ng dramatically affects the properties of this integrator.

## 4 Discussion

CaMKII phosphorylation is a fundamental process downstream of *Ca*^2+^-influx in the PSD. There have been many studies focused on the interactions of *Ca*^2+^, CaM, and CaMKII, and their role in the dynamics of CaMKII phosphorylation [74, 77, 43, 94, 59, 72, 85]. In the work presented here, using computational modeling, we show that the presence of a scaffolding molecule, Ng, has a non-linear effect on the dose response curve of CaMKII phosphorylation in response to *Ca*^2+^ influx. Our results show that CaMKII holoenzyme is less sensitive to *Ca*^2+^ signals than the equivalent number of monomers, and thus less susceptible to noise and fluctuations of [*Ca*^2+^]. Ng further modulates CaMKII phosphorylation in response to [*Ca*^2+^] spikes in the physiological range of [CaM]. Finally, we predict that in the presence of PP1, the CaMKII holoenzyme acts as a leaky integrator of calcium signals, and Ng significantly affects both the capacity and the leak rate of this integrator (Figure 9C). Based on these findings, we conclude that Ng plays a crucial role in fine-tuning the postsynaptic response to *Ca*^2+^ signals.

This prediction is consistent with experimental data that show that Ng knockout mice exhibit a decrease in LTP induction and spatial learning [71]. Experimental results show that impairment of the binding of Ng to CaM results in lower activity of CaMKII in the dendritic spines [49, 71]. At first sight, this finding seems contradictory to what we find here: the presence of Ng decreases CaMKII activation in our simulations. To understand this apparent inconsistency we note that in our simulations, we compared the results obtained with and without Ng while keeping the overall [CaM] the same for both cases. *In vivo* however, the scaffolding protein Ng sequesters CaM into the dendritic spines, thereby increasing the overall [CaM] in the spines that can be released upon stimulation [27]. This explanation is supported by the observation that postsynaptic injection of *Ca*^2+^*/CaM* enhances the synaptic strength in the same manner as overexpression of neurogranin in CA1 neurons [27, 102, 95].

The role of Ng in modulating the dynamics of CaMKII, particularly, the leak rate and capacity of CaMKII phosphorylation have multiple implications in downstream processes in a spine. Phosphorylation of CaMKII activates its kinase domain, which leads to subsequent modification of the dendritic spine and the postsynaptic density [52, 38, 24]. Specifically phosphorylated CaMKII unbinds from F-actin in the spine allowing reorganization of the cytoskeleton, and rebinds this newly structured F-actin after it has been dephosphorylated [84, 99, 30, 93, 18, 38]. These interactions with the actin cytoskeleton allow for dynamic rearrangements of the actin organization within the spine, subsequently impacting its size and shape [41, 45, 83] (Figure 10). CaMKII activation also leads to increased trafficking [103, 70] and trapping [1] of AMPA receptors at the PSD as well as an increase in the conductance of these receptors after the induction of LTP [4, 8, 51, 48]. All of these events result in changes to the size and strength of the synapse [78, 55, 2, 3] (Figure 10).

**Figure 10:**
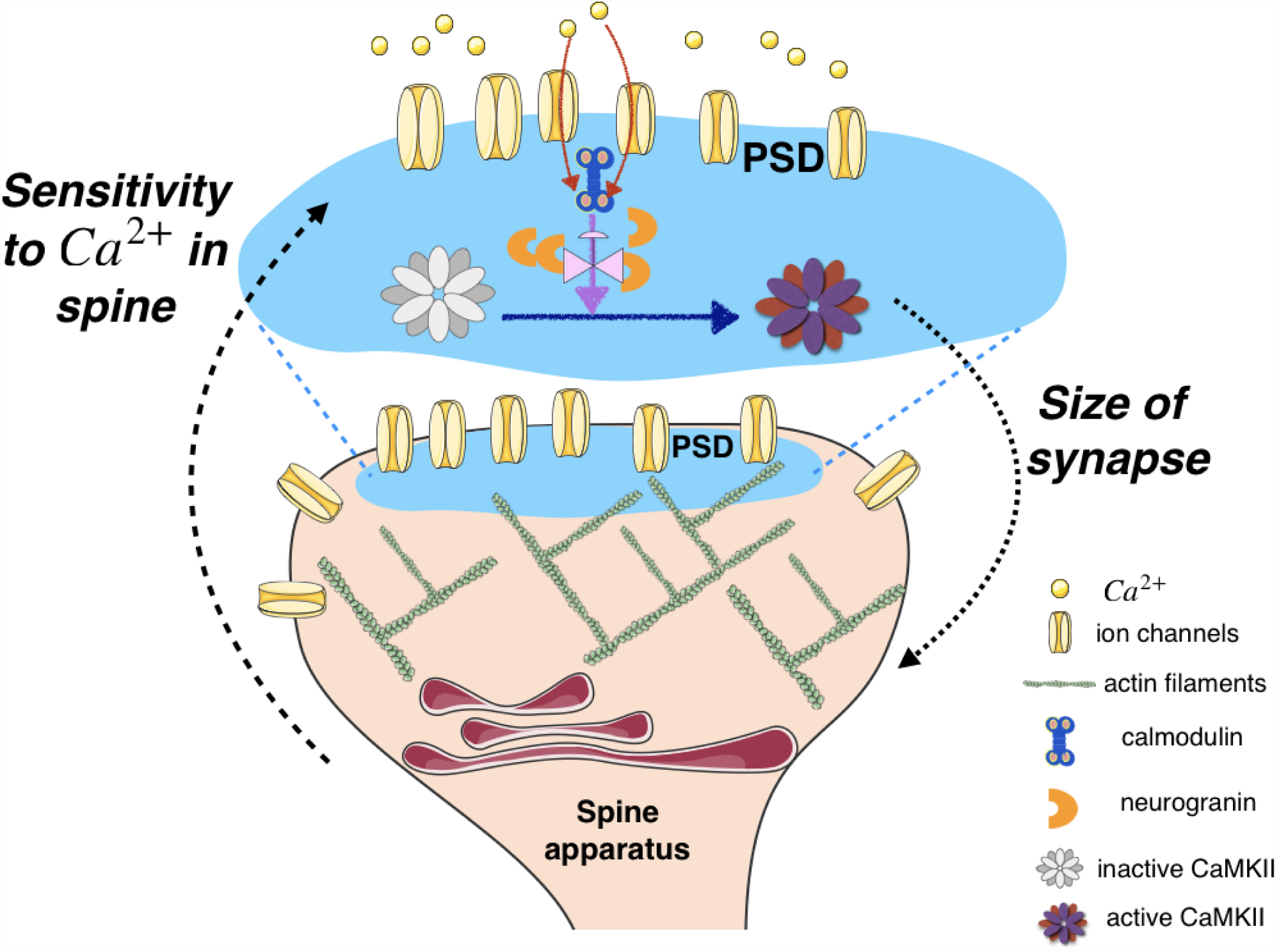
Schematic representation of CaMKII phosphorylation controlling sensitivity of a synapse to *Ca*^2+^ signals. In the PSD, influx of *Ca*^2+^ through the ion channel, binding of *Ca*^2+^ to calmodulin, and phosphorylation of CaMKII sets off the series of events associated with synaptic plasticity. In this work, we show that the competition between neurogranin sequestration of calmodulin and *Ca*^2+^ binding of calmodulin affects CaMKII dynamics. This has implications for the feedback between *Ca*^2+^ in the spine and structural plasticity of the spine, particularly size dynamics.

Synaptic size and strength are regulated by firing history [11] with very high precision [6]. As a synapse undergoes increased LTP, it grows larger and subsequently the *Ca*^2+^ transients in the spine become increasingly dilute in response to back-propagating action potentials [5]. Thus, the integration of *Ca*^2+^ signals is an built-in mechanism in the dendritic spine to down-regulate its sensitivity to these signals as the corresponding synapse grows larger (Figure 10) and our study shows that scaffolding molecules such as Ng can modulate these dynamics. The results of our simulations combined with experimental findings indicated above point to the extreme importance of Ng in regulating the postsynaptic response to *Ca*^2+^ signals. This finding adds to the growing evidence that scaffolding proteins fine-tune the signaling pathways in biological signal transduction mechanisms [10, 47, 56].

The model described here will enable the development of more detailed models of signaling events in the PSD including coupling between CaMKII and PKA [66, 69, 88], spatial organization of calcium-CaM-CaMKII dynamics building on [7, 67, 25, 6, 5], and coupling between CaMKII and actin interactions [45, 46]. Such efforts will be necessary to inform the mechanochemical coupling within spines and how they regulate information processing.

## 5 Acknowledgments

We thank Miriam Bell for useful discussions and assistance with making figure 10. We also thank Donya Ohadi and Andrea Cugno for useful discussions. This work was supported by the following grants: FA9550-18-1-0051 (PR,MO,TB,TS), NIH: P41GM103712 (TS,TB), NS44306 (MK,TB,TS), DA030749 (MK,TB,TS).

## References

[1] V. J. Appleby, S. A. Corrêa, J. K. Duckworth, J. E. Nash, J. Noël, S. M. Fitzjohn, G. L. Collingridge, and E. Molnár. Ltp in hippocampal neurons is associated with a camkii-mediated increase in glua1 surface expression. Journal of neurochemistry, 116(4):530–543, 2011.

[2] B. Asrican, J. Lisman, and N. Otmakhov. Synaptic strength of individual spines correlates with bound ca2+–calmodulin-dependent kinase ii. Journal of Neuroscience, 27(51):14007–14011, 2007.

[3] K. Barcomb, J. W. Hell, T. A. Benke, and K. U. Bayer. The camkii/glun2b protein interaction maintains synaptic strength. Journal of Biological Chemistry, 291(31):16082–16089, 2016.

[4] A. Barria, D. Muller, V. Derkach, L. C. Griffith, and T. R. Soderling. Regulatory phosphorylation of ampa-type glutamate receptors by cam-kii during long-term potentiation. Science, 276(5321):2042–2045, 1997.

[5] T. M. Bartol, D. X. Keller, J. P. Kinney, C. L. Bajaj, K. M. Harris, T. J. Sejnowski, and M. B. Kennedy. Computational reconstitution of spine calcium transients from individual proteins. Frontiers in synaptic neuroscience, 7:17, 2015.

[6] T. M. Bartol Jr, C. Bromer, J. Kinney, M. A. Chirillo, J. N. Bourne, K. M. Harris, and T. J. Sejnowski. Nanoconnectomic upper bound on the variability of synaptic plasticity. Elife, 4:e10778, 2015.

[7] M. Bell, T. Bartol, T. Sejnowski, and P. Rangamani. Addendum: Dendritic spine geometry and spine apparatus organization govern the spatiotemporal dynamics of calcium. The Journal of general physiology, 151(9):2221, 2019.

[8] T. A. Benke, A. Lüthi, J. T. Isaac, and G. L. Collingridge. Modulation of ampa receptor unitary conductance by synaptic activity. Nature, 393(6687):793, 1998.

[9] M. K. Bennett, N. E. Erondu, and M. B. Kennedy. Purification and characterization of a calmodulin-dependent protein kinase that is highly concentrated in brain. Journal of Biological Chemistry, 258(20):12735–12744, 1983.

[10] R. P. Bhattacharyya, A. Reményi, M. C. Good, C. J. Bashor, A. M. Falick, and W. A. Lim. The ste5 scaffold allosterically modulates signaling output of the yeast mating pathway. Science, 311(5762):822–826, 2006.

[11] G.-q. Bi and M.-m. Poo. Synaptic modifications in cultured hippocampal neurons: dependence on spike timing, synaptic strength, and postsynaptic cell type. Journal of neuroscience, 18(24):10464–10472, 1998.

[12] A. Biber, G. Schmid, and K. Hempel. Calmodulin content in specific brain areas. Experimental brain research, 56(2):323–326, 1984.

[13] M. L. Blinov, J. R. Faeder, B. Goldstein, and W. S. Hlavacek. Bionetgen: software for rule-based modeling of signal transduction based on the interactions of molecular domains. Bioinformatics, 20(17):3289–3291, 2004.

[14] J. Borovac, M. Bosch, and K. Okamoto. Regulation of actin dynamics during structural plasticity of dendritic spines: Signaling messengers and actin-binding proteins. Molecular and Cellular Neuroscience, 2018.

[15] J. M. Bradshaw, A. Hudmon, and H. Schulman. Chemical quenched flow kinetic studies indicate an intraholoenzyme autophosphorylation mechanism for ca2+/calmodulin-dependent protein kinase ii. Journal of Biological Chemistry, 277(23):20991–20998, 2002.

[16] J. M. Bradshaw, Y. Kubota, T. Meyer, and H. Schulman. An ultrasensitive ca2+/calmodulin-dependent protein kinase ii-protein phosphatase 1 switch facilitates specificity in postsynaptic calcium signaling. Proceedings of the National Academy of Sciences, 100(18):10512–10517, 2003.

[17] S. E. Brown, S. R. Martin, and P. M. Bayley. Kinetic control of the dissociation pathway of calmodulin-peptide complexes. Journal of Biological Chemistry, 272(6):3389–3397, 1997.

[18] H. J. Carlisle, P. Manzerra, E. Marcora, and M. B. Kennedy. Syngap regulates steady-state and activity-dependent phosphorylation of cofilin. Journal of Neuroscience, 28(50):13673–13683, 2008.

[19] J.-Y. Chang, P. Parra-Bueno, T. Laviv, E. M. Szatmari, S.-J. R. Lee, and R. Yasuda. Camkii autophosphorylation is necessary for optimal integration of ca 2+ signals during ltp induction, but not maintenance. Neuron, 94(4):800–808, 2017.

[20] L. H. Chao, M. M. Stratton, I.-H. Lee, O. S. Rosenberg, J. Levitz, D. J. Mandell, T. Kortemme, J. T. Groves, H. Schulman, and J. Kuriyan. A mechanism for tunable autoinhibition in the structure of a human ca2+/calmodulin-dependent kinase ii holoenzyme. Cell, 146(5):732–745, 2011.

[21] R. J. Colbran. Protein phosphatases and calcium/calmodulin-dependent protein kinase ii-dependent synaptic plasticity. Journal of Neuroscience, 24(39):8404–8409, 2004.

[22] R. A. Copeland. Enzymes: a practical introduction to structure, mechanism, and data analysis. John Wiley & Sons, 2004.

[23] S. J. Coultrap and K. U. Bayer. Camkii regulation in information processing and storage. Trends in neurosciences, 35(10):607–618, 2012.

[24] S. J. Coultrap, R. K. Freund, H. O’Leary, J. L. Sanderson, K. W. Roche, M. L. Dell’Acqua, and K. U. Bayer. Autonomous camkii mediates both ltp and ltd using a mechanism for differential substrate site selection. Cell reports, 6(3):431–437, 2014.

[25] A. Cugno, T. M. Bartol, T. J. Sejnowski, R. Iyengar, and P. Rangamani. Geometric principles of second messenger dynamics in dendritic spines. Scientific reports, 9(1):1–18, 2019.

[26] W. Denk, R. Yuste, K. Svoboda, and D. W. Tank. Imaging calcium dynamics in dendritic spines. Current opinion in neurobiology, 6(3):372–378, 1996.

[27] F. J. Díez-Guerra. Neurogranin, a link between calcium/calmodulin and protein kinase c signaling in synaptic plasticity. IUBMB life, 62(8):597–606, 2010.

[28] J. R. Faeder, M. L. Blinov, and W. S. Hlavacek. Rule-based modeling of biochemical systems with 26 bionetgen. In Systems biology, pages 113–167. Springer, 2009.

[29] B. E. Finn and S. Forsén. The evolving model of calmodulin structure, function and activation. Structure, 3(1):7–11, 1995.

[30] I. N. Fleming, C. M. Elliott, and J. H. Exton. Phospholipase c-*γ*, protein kinase c and ca2+/calmodulin-dependent protein kinase ii are involved in platelet-derived growth factor-induced phosphorylation of tiam1. FEBS letters, 429(3):229–233, 1998.

[31] K. M. Franks and T. J. Sejnowski. Complexity of calcium signaling in synaptic spines. Bioessays, 24(12):1130–1144, 2002.

[32] H. Fujii, M. Inoue, H. Okuno, Y. Sano, S. Takemoto-Kimura, K. Kitamura, M. Kano, and H. Bito. Nonlinear decoding and asymmetric representation of neuronal input information by camkii*α* and calcineurin. Cell reports, 3(4):978–987, 2013.

[33] T. R. Gaertner, J. A. Putkey, and M. N. Waxham. Rc3/neurogranin and ca2+/calmodulin-dependent protein kinase ii produce opposing effects on the affinity of calmodulin for calcium. Journal of Biological Chemistry, 279(38):39374–39382, 2004.

[34] P. I. Hanson, M. S. Kapiloff, L. L. Lou, M. G. Rosenfeld, and H. Schulman. Expression of a multifunctional ca2+/calmodulin-dependent protein kinase and mutational analysis of its autoregulation. Neuron, 3(1):59–70, 1989.

[35] P. I. Hanson, T. Meyer, L. Stryer, and H. Schulman. Dual role of calmodulin in autophosphorylation of multifunctional cam kinase may underlie decoding of calcium signals. Neuron, 12(5):943–956, 1994.

[36] P. I. Hanson and H. Schulman. Inhibitory autophosphorylation of multifunctional ca2+/calmodulin-dependent protein kinase analyzed by site-directed mutagenesis. Journal of Biological Chemistry, 267(24):17216–17224, 1992.

[37] L. A. Harris, J. S. Hogg, J.-J. Tapia, J. A. Sekar, S. Gupta, I. Korsunsky, A. Arora, D. Barua, R. P. Sheehan, and J. R. Faeder. Bionetgen 2.2: advances in rule-based modeling. Bioinformatics, 32(21):3366–3368, 2016.

[38] Y. Hayashi, S.-H. Shi, J. A. Esteban, A. Piccini, J.-C. Poncer, and R. Malinow. Driving ampa receptors into synapses by ltp and camkii: requirement for glur1 and pdz domain interaction. Science, 287(5461):2262–2267, 2000.

[39] M. J. Higley and B. L. Sabatini. Calcium signaling in dendritic spines. Cold Spring Harbor perspectives in biology, 4(4):a005686, 2012.

[40] L. Hoffman, A. Chandrasekar, X. Wang, J. A. Putkey, and M. N. Waxham. Neurogranin alters the structure and calcium-binding properties of calmodulin. Journal of Biological Chemistry, pages jbc–M114, 2014.

[41] L. Hoffman, M. M. Farley, and M. N. Waxham. Calcium-calmodulin-dependent protein kinase ii isoforms differentially impact the dynamics and structure of the actin cytoskeleton. Biochemistry, 52(7):1198–1207, 2013.

[42] A. Hudmon and H. Schulman. Neuronal ca2+/calmodulin-dependent protein kinase ii: the role of structure and autoregulation in cellular function. Annual review of biochemistry, 71(1):473–510, 2002.

[43] T. Johnson, T. Bartol, T. Sejnowski, and E. Mjolsness. Model reduction for stochastic camkii reaction kinetics in synapses by graph-constrained correlation dynamics. Physical biology, 12(4):045005, 2015.

[44] S. Kakiuchi, S. Yasuda, R. Yamazaki, Y. Teshima, K. Kanda, R. Kakiuchi, and K. Sobue. Quantitative determinations of calmodulin in the supernatant and particulate fractions of mammalian tissues. The Journal of Biochemistry, 92(4):1041–1048, 1982.

[45] S. Khan, I. Conte, T. Carter, K. U. Bayer, and J. E. Molloy. Multiple camkii binding modes to the actin cytoskeleton revealed by single-molecule imaging. Biophysical journal, 111(2):395–408, 2016.

[46] S. Khan, K. H. Downing, and J. E. Molloy. Architectural dynamics of camkii-actin networks. Biophysical journal, 116(1):104–119, 2019.

[47] W. Kolch. Coordinating erk/mapk signalling through scaffolds and inhibitors. Nature reviews Molecular cell biology, 6(11):827, 2005.

[48] G. Krapivinsky, I. Medina, L. Krapivinsky, S. Gapon, and D. E. Clapham. Syngap-mupp1-camkii synaptic complexes regulate p38 map kinase activity and nmda receptor-dependent synaptic ampa receptor potentiation. Neuron, 43(4):563–574, 2004.

[49] T. Krucker, G. R. Siggins, R. K. McNamara, K. A. Lindsley, A. Dao, D. W. Allison, L. De Lecea, T. W. Lovenberg, J. G. Sutcliffe, and D. D. Gerendasy. Targeted disruption of rc3 reveals a calmodulin-based mechanism for regulating metaplasticity in the hippocampus. Journal of Neuroscience, 22(13):5525–5535, 2002.

[50] Y. Kubota, J. A. Putkey, and M. N. Waxham. Neurogranin controls the spatiotemporal pattern of postsynaptic ca2+/cam signaling. Biophysical journal, 93(11):3848–3859, 2007.

[51] H.-K. Lee, M. Barbarosie, K. Kameyama, M. F. Bear, and R. L. Huganir. Regulation of distinct ampa receptor phosphorylation sites during bidirectional synaptic plasticity. Nature, 405(6789):955, 2000.

[52] S.-J. R. Lee, Y. Escobedo-Lozoya, E. M. Szatmari, and R. Yasuda. Activation of camkii in single dendritic spines during long-term potentiation. Nature, 458(7236):299, 2009.

[53] S. Linse, A. Helmersson, and S. Forsen. Calcium binding to calmodulin and its globular domains. Journal of Biological Chemistry, 266(13):8050–8054, 1991.

[54] J. Lisman, H. Schulman, and H. Cline. The molecular basis of camkii function in synaptic and behavioural memory. Nature Reviews Neuroscience, 3(3):175, 2002.

[55] J. Lisman, R. Yasuda, and S. Raghavachari. Mechanisms of camkii action in long-term potentiation. Nature reviews neuroscience, 13(3):169, 2012.

[56] J. W. Locasale, A. S. Shaw, and A. K. Chakraborty. Scaffold proteins confer diverse regulatory properties to protein kinase cascades. Proceedings of the National Academy of Sciences, 104(33):13307–13312, 2007.

[57] S. R. Martin, J. F. Maune, K. Beckingham, and P. M. Bayley. Stopped-flow studies of calcium dissociation from calcium-binding-site mutants of drosophila melanogaster calmodulin. European journal of biochemistry, 205(3):1107–1117, 1992.

[58] T. Meyer, P. I. Hanson, L. Stryer, and H. Schulman. Calmodulin trapping by calcium-calmodulin-dependent protein kinase. Science, 256(5060):1199–1202, 1992.

[59] S. G. Miller and M. B. Kennedy. Regulation of brain type ii ca2+ calmodulin-dependent protein kinase by autophosphorylation: A ca2+-triggered molecular switch. Cell, 44(6):861–870, 1986.

[60] S. G. Miller, B. L. Patton, and M. B. Kennedy. Sequences of autophosphorylation sites in neuronal type ii cam kinase that control ca2+-independent activity. Neuron, 1(7):593–604, 1988.

[61] V. N. Murthy, T. Schikorski, C. F. Stevens, and Y. Zhu. Inactivity produces increases in neurotransmitter release and synapse size. Neuron, 32(4):673–682, 2001.

[62] J. B. Myers, V. Zaegel, S. J. Coultrap, A. P. Miller, K. U. Bayer, and S. L. Reichow. The camkii holoenzyme structure in activation-competent conformations. Nature communications, 8:15742, 2017.

[63] E. Neher and G. Augustine. Calcium gradients and buffers in bovine chromaffin cells. The Journal of physiology, 450(1):273–301, 1992.

[64] T. Nevian and B. Sakmann. Spine ca2+ signaling in spike-timing-dependent plasticity. Journal of Neuroscience, 26(43):11001–11013, 2006.

[65] E. A. Nimchinsky, R. Yasuda, T. G. Oertner, and K. Svoboda. The number of glutamate receptors opened by synaptic stimulation in single hippocampal spines. Journal of Neuroscience, 24(8):2054–2064, 2004.

[66] D. Ohadi and P. Rangamani. Geometric control of frequency modulation of camp oscillations due to ca2+-bursts in dendritic spines. bioRxiv, page 520643, 2019.

[67] D. Ohadi, D. L. Schmitt, B. Calabrese, S. Halpain, J. Zhang, and P. Rangamani. Computational modeling reveals frequency modulation of calcium-camp/pka pathway in dendritic spines. bioRxiv, page 521740, 2019.

[68] K.-I. Okamoto, R. Narayanan, S. H. Lee, K. Murata, and Y. Hayashi. The role of camkii as an f-actin-bundling protein crucial for maintenance of dendritic spine structure. Proceedings of the National Academy of Sciences, 104(15):6418–6423, 2007.

[69] R. F. Oliveira, A. Terrin, G. Di Benedetto, R. C. Cannon, W. Koh, M. Kim, M. Zaccolo, and K. T. Blackwell. The role of type 4 phosphodiesterases in generating microdomains of camp: large scale stochastic simulations. PloS one, 5(7):e11725, 2010.

[70] P. Opazo and D. Choquet. A three-step model for the synaptic recruitment of ampa receptors. Molecular and Cellular Neuroscience, 46(1):1–8, 2011.

[71] J. H. Pak, F. L. Huang, J. Li, D. Balschun, K. G. Reymann, C. Chiang, H. Westphal, and K.-P. Huang. Involvement of neurogranin in the modulation of calcium/calmodulin-dependent protein kinase ii, synaptic plasticity, and spatial learning: a study with knockout mice. Proceedings of the National Academy of Sciences, 97(21):11232–11237, 2000.

[72] B. L. Patton, S. G. Miller, and M. B. Kennedy. Activation of type ii calcium/calmodulin-dependent protein kinase by ca2+/calmodulin is inhibited by autophosphorylation of threonine within the calmodulin-binding domain. Journal of Biological Chemistry, 265(19):11204–11212, 1990.

[73] O. B. Peersen, T. S. Madsen, and J. J. Falke. Intermolecular tuning of calmodulin by target peptides and proteins: differential effects on ca2+ binding and implications for kinase activation. Protein Science, 6(4):794–807, 1997.

[74] S. Pepke, T. Kinzer-Ursem, S. Mihalas, and M. B. Kennedy. A dynamic model of interactions of ca2+, calmodulin, and catalytic subunits of ca2+/calmodulin-dependent protein kinase ii. PLoS computational biology, 6(2):e1000675, 2010.

[75] A. Persechini, H. D. White, and K. J. Gansz. Different mechanisms for ca dissociation from complexes of calmodulin with nitric oxide synthase or myosin light chain kinase. Journal of Biological Chemistry, 271(1):62–67, 1996.

[76] J. J. Petrozzino, L. D. P. Miller, and J. A. Connor. Micromolar ca2+ transients in dendritic spines of hippocampal pyramidal neurons in brain slice. Neuron, 14(6):1223–1231, 1995.

[77] M. C. Pharris, T. M. Bartol, T. J. Sejnowski, M. B. Kennedy, M. I. Stefan, and T. L. Kinzer-Ursem. A multi-state model of the camkii dodecamer suggests a role for calmodulin in maintenance of autophosphorylation. 2019.

[78] H. J. Pi, N. Otmakhov, F. El Gaamouch, D. Lemelin, P. De Koninck, and J. Lisman. Camkii control of spine size and synaptic strength: role of phosphorylation states and nonenzymatic action. Proceedings of the National Academy of Sciences, 107(32):14437–14442, 2010.

[79] P. Rangamani, M. G. Levy, S. Khan, and G. Oster. Paradoxical signaling regulates structural plasticity in dendritic spines. Proceedings of the National Academy of Sciences, 113(36):E5298–E5307, 2016.

[80] R. C. Rich and H. Schulman. Substrate-directed function of calmodulin in autophosphorylation of ca2+/calmodulin-dependent protein kinase ii. Journal of Biological Chemistry, 273(43):28424–28429, 1998.

[81] O. S. Rosenberg, S. Deindl, R.-J. Sung, A. C. Nairn, and J. Kuriyan. Structure of the autoinhibited kinase domain of camkii and saxs analysis of the holoenzyme. Cell, 123(5):849–860, 2005.

[82] B. L. Sabatini, T. G. Oertner, and K. Svoboda. The life cycle of ca2+ ions in dendritic spines. Neuron, 33(3):439–452, 2002.

[83] H. Sanabria, M. T. Swulius, S. J. Kolodziej, J. Liu, and M. N. Waxham. *β*camkii regulates actin assembly and structure. Journal of Biological Chemistry, 284(15):9770–9780, 2009.

[84] K. Shen, M. N. Teruel, K. Subramanian, and T. Meyer. Camkii*β* functions as an f-actin targeting module that localizes camkii*α*/*β* heterooligomers to dendritic spines. Neuron, 21(3):593–606, 1998.

[85] J. M. Shifman, M. H. Choi, S. Mihalas, S. L. Mayo, and M. B. Kennedy. Ca2+/calmodulin-dependent protein kinase ii (camkii) is activated by calmodulin with two bound calciums. Proceedings of the National Academy of Sciences, 103(38):13968–13973, 2006.

[86] A. J. Silva, R. Paylor, J. M. Wehner, and S. Tonegawa. Impaired spatial learning in alpha-calcium-calmodulin kinase ii mutant mice. Science, 257(5067):206–211, 1992.

[87] A. J. Silva, C. F. Stevens, S. Tonegawa, and Y. Wang. Deficient hippocampal long-term potentiation in alpha-calcium-calmodulin kinase ii mutant mice. Science, 257(5067):201–206, 1992.

[88] D. Singh and U. S. Bhalla. Subunit exchange enhances information retention by camkii in dendritic spines. Elife, 7:e41412, 2018.

[89] M. W. Sneddon, J. R. Faeder, and T. Emonet. Efficient modeling, simulation and coarse-graining of biological complexity with nfsim. Nature methods, 8(2):177, 2011.

[90] S. Strack, S. Choi, D. M. Lovinger, and R. J. Colbran. Translocation of autophosphorylated calcium/calmodulin-dependent protein kinase ii to the postsynaptic density. Journal of Biological Chemistry, 272(21):13467–13470, 1997.

[91] K. Takao, K.-I. Okamoto, T. Nakagawa, R. L. Neve, T. Nagai, A. Miyawaki, T. Hashikawa, S. Kobayashi, and Y. Hayashi. Visualization of synaptic ca2+/calmodulin-dependent protein kinase ii activity in living neurons. Journal of Neuroscience, 25(12):3107–3112, 2005.

[92] J.-J. Tapia, A. S. Saglam, J. Czech, R. Kuczewski, T. M. Bartol, T. J. Sejnowski, and J. R. Faeder. Mcell-r: A particle-resolution network-free spatial modeling framework. In Modeling Biomolecular Site Dynamics, pages 203–229. Springer, 2019.

[93] K. F. Tolias, J. B. Bikoff, A. Burette, S. Paradis, D. Harrar, S. Tavazoie, R. J. Weinberg, and M. E. Greenberg. The rac1-gef tiam1 couples the nmda receptor to the activity-dependent development of dendritic arbors and spines. Neuron, 45(4):525–538, 2005.

[94] J. K. Tse, A. M. Giannetti, and J. M. Bradshaw. Thermodynamics of calmodulin trapping by ca2+/calmodulin-dependent protein kinase ii: Subpicomolar k d determined using competition titration calorimetry. Biochemistry, 46(13):4017–4027, 2007.

[95] J.-H. Wang and P. T. Kelly. Postsynaptic injection of ca2+/cam induces synaptic potentiation requiring camkii and pkc activity. Neuron, 15(2):443–452, 1995.

[96] D. Watterson, W. Harrelson, P. Keller, F. Sharief, and T. C. Vanaman. Structural similarities between the ca2+-dependent regulatory proteins of 3’: 5’-cyclic nucleotide phosphodiesterase and actomyosin atpase. Journal of Biological Chemistry, 251(15):4501–4513, 1976.

[97] G.-Y. Wu and H. T. Cline. Stabilization of dendritic arbor structure in vivo by camkii. Science, 279(5348):222–226, 1998.

[98] Z. Xia and D. R. Storm. The role of calmodulin as a signal integrator for synaptic plasticity. Nature Reviews Neuroscience, 6(4):267, 2005.

[99] Z. Xie, D. P. Srivastava, H. Photowala, L. Kai, M. E. Cahill, K. M. Woolfrey, C. Y. Shum, D. J. Surmeier, and P. Penzes. Kalirin-7 controls activity-dependent structural and functional plasticity of dendritic spines. Neuron, 56(4):640–656, 2007.

[100] S.-N. Yang, Y.-G. Tang, and R. S. Zucker. Selective induction of ltp and ltd by postsynaptic [ca2+] i elevation. Journal of neurophysiology, 81(2):781–787, 1999.

[101] A. M. Zhabotinsky, R. N. Camp, I. R. Epstein, and J. E. Lisman. Role of the neurogranin concentrated in spines in the induction of long-term potentiation. Journal of Neuroscience, 26(28):7337–7347, 2006.

[102] L. Zhong, T. Cherry, C. E. Bies, M. A. Florence, and N. Z. Gerges. Neurogranin enhances synaptic strength through its interaction with calmodulin. The EMBO journal, 28(19):3027–3039, 2009.

[103] J. J. Zhu, Y. Qin, M. Zhao, L. Van Aelst, and R. Malinow. Ras and rap control ampa receptor trafficking during synaptic plasticity. Cell, 110(4):443–455, 2002.

[104] D.-J. Zou and H. T. Cline. Postsynaptic calcium/calmodulin-dependent protein kinase ii is required to limit elaboration of presynaptic and postsynaptic neuronal arbors. Journal of Neuroscience, 19(20):8909–8918, 1999.

